# Expression and novel alternative purification of the recombinant nucleocapsid (N) protein of SARS-CoV-2 in Escherichia coli for the serodiagnosis of COVID-19

**DOI:** 10.1101/2021.11.10.467990

**Authors:** Jose David Rosales, William Quintero, Jhon Cruz, Balbino Perdomo, Militza Quintero, Marcos Bastidas, Jose Domingo Lugo, Keila Rivas Rodriguez, Juan Carlos Freites-Perez, Annie Castillo

## Abstract

The SARS-CoV-2 coronavirus causes severe acute respiratory syndrome and has caused a global pandemic by causing the COVID-19 disease. To monitor and control it, diagnostic methods such as molecular and serological tests are necessary. The serological approach uses SARS-CoV-2 antigens to detect the antibodies present in patients using quantitative techniques such as enzyme-linked immunosorbent assay (ELISA) or qualitative rapid tests such as lateral flow chromatography (RDT’s). The main antigens used are the spike protein (S) and the nucleocapsid protein (N). Both proteins are obtained in different expression systems, in eukaryotic cells, their production is expensive, so in this work we chose a simpler and cheaper system such as prokaryotic to express and purify the N protein. Thereore, the nucleotide sequence had to being optimized to be expressed in Escherichia coli. The protein N is sensitive to *E.coli* proteases and also has the ability to self-proteolyze under native conditions, degrading into different fragments. However, under denaturing conditions, using urea and at pH 5.3 it is stable and efficiently purified using metal exchange chromatography (IMAC). In our purification strategy, we surprisingly found that by not using a sonicator, a homogeneous and time-stable preparation of the recombinant antigen is obtained. An approximate yield of 200 mg / L was obtained. It was then tested with healthy sera and sera from COVID-19 convalescent patients in Wester-blot tests that were able to recognize it. Our work provides a novel strategy to produce the SARS-CoV-2 protein N so that it can be used as an input in the development and innovation of serological tests in the diagnosis of COVID-19.

## Introduction

The etiological agent of the severe acute respiratory syndrome (SARS-CoV-2) that emerged in 2019 has caused 250 million cases and 5 million deaths until November 2021 (WHO, 2021), being identified as a new type of coronavirus with a genome of single-stranded positive sense RNA (+ ssRNA) of 29,903 nucleotides, and its genomic sequence being reported for the first time in March 2020 (1), Genome ID: MN908947.3. Since then, thousands of genomes have been sequenced and potentially more contagious and serious new viral variants have been identified such as: Alpha (B.1.1.7), Beta (B.1.351, Gamma (P.1), Epsilon (B.1.427 and B.1.429), Eta (B.1.525), Iota (B.1.526), Kappa (B.1.617.1), Mu (B.1.621, B.1.621.1), Zeta (P.2) and delta (B.1.617) according to the Center for Disease Control and Prevention (5).

RNA detection is performed by reverse transcription polymerase chain reaction (RT-PCR), although there are other very promising molecular diagnostic methods including those based on Regularly Inter-spaced Clustered Short Palindromic Repeats (CRISPR), reverse transcription loop-mediated isothermal amplification (RT-LAMP), digital PCR, microarray assays, and next-generation sequencing (NGS) (3).

Molecular techniques are highly sensitive, consume a lot of resources, time and in certain conditions, where patients suffer from heart disease, immunosuppression and diabetes, a false negative result could be obtain (4), for which serological and clinical methods have been used to complement the analysis and help in containing the pandemic (5–6).

Serological tests provide rapid results by detecting antibodies against SARS-CoV-2 in blsood, plasma or serum sample in most infected patients in the first 15 days after the start of the clinic and up to 3 months after infection, although, this time will depend on the immune response of each person.

This means that there is a window period where molecular techniques will stop detecting the presence of SARS-CoV-2 and will give negative PCR, while the anti-SARS-CoV-2 IgG antibodies will remain and give positive results (7). These tests will depend on the selected antigen, such as spike protein (S) and nucleocapsid (N) (8–9). Having recombinant antigens for serological diagnosis makes it possible to analyze the immune response to SARS-CoV-2, detect and characterize specific antibodies, and carry out large-scale epidemiological studies within the population. These serological tests will also help to evaluate the efficacy of the different drugs and vaccines available (10).

Protein N is a protein is made up of 419 amino acids, with two domains rich in arginine and lysine, whose molecular weight varies between 45 and 60 kDa, depending on its post-translational modifications (11),and it plays an important role in the process of assembly of particles of virus, enveloping complete genomic RNA (12). Protein N is a powerful antigen for the diagnosis of SARS-CoV-2 infection by detecting IgM and IgG antibodies and is used in different immunological techniques such as chemiluminescent microparticle immunoassay (CMIA) (13), ELISA (14), Wester-blot (15) and rapid tests (16). Recombinant protein N is obtained from *E.coli*, baculovirus and mammalian cells (17–20). In our work we provide a simple strategy to produce SARS-CoV-2 Protein N in *Escherichia coli* and we demonstrate that it is recognized by positive sera from COVID-19 patients.

## Materials and methods

### Synthetic construction N-pET20b

The sequence of the SARS-CoV-2 recombinant protein N gene is obtained from the GenBank accession: MN908947.3 region 28274-29533. The optimized sequence is obtained synthetically and cloned into the prokaryotic expression vector pET20b (GenScript, USA). The promoter T7 oligos were used: AATACGACTCACTATAGG and terminator T7: GCTAGTTATTGCTCAGCGG, and a DNA polymerase (Promega., USA) for PCR, using the following parameters: heating at 95 ° C for 5 min; 32 cycles of denaturation at 95 ° C for 1 min, hybridization at 45 ° C for 1 min and extension at 72 ° C for 2 min.

### Culture media

Peptone, casein, yeast extract, NaCl (Promega, USA). Isopropyl-β-D-thiogalactopyranoside (IPTG), anti-human IgG conjugate (SIGMA-ALDRID, USA). Ni-NTA agarose resin (Invitrogen, USA).

### Transformation and verification of the expression vector in E. coli

The plasmid pET20b-N (4ug) is resuspended in 100 uL of Megapure water and 5 ng is used to transform the *E. coli* Top10 (Invitrogen, USA) to maintain the plasmid and in the strain of *E. coli* BL21 (DE3) (Novagen, USA) for your expression. Briefly, the transformation was carried out with the CaCl_2_ chemical method and was placed in an LB plate containing 100 μg / ml of ampicillin and incubated at 37°C in a 12-16 h incubator. A PCR is performed from a colony to verify the insert. The fragment was visualized and identified by 1% DNA agarose gel electrophoresis.

### Expression of protein N in *E. coli* cepa BL21 (DE3)

The construction of the recombinant SARS-CoV-2 N protein in the pET20b vector encodes a protein of 667 amino acids with a weight of 51 kDa. Induction was carried out in LB medium at 18° C with 1mM IPTG. Bacteria were collected by centrifugation (12000 g for 15 min at 4° C) and frozen at −20°C until use. Expression was verified on 15% SDS-PAGE gels, according to Laemmli (21). When verifying the soluble and insoluble fractions, all the protein was found forming inclusion bodies.

### Recombinant nucleocapsid purification

#### Hybrid Conditions

Bacterial cells were resuspended in a solution consisting of Tris (100 mM) pH 8.0, NaCl (300 mM) and lysozyme (1 mg / ml), PMSF (1mM) and 10 uL of a protease inhibitor cocktail. It was incubated at room temperature for 60 min and frozen for 16 h, thawed and the pellet was separated from the supernatant by centrifugation at 12000 rpm. Two conditions were used. In the first (hybrid condition) the pellet was resuspended in 100 mM phosphate buffer (pH 8.0), 300 NaCl, 10mM Imidazole and 8M guanidine. It was incubated for 16 h with shaking and centrifuged at 12000 rpm. The supernatant was used to load a column with IMAC ProBond ™ resin (Thermofisher, USA) and equilibrated with the same buffer. 5 column volumes were washed with PBS with 10 mM imidazole and 8M guanidine, to remove non-specific proteins. Then it was washed with 5 column volumes with PBS without guanidine. The protein is then eluted with PBS containing 250 mM imidazole. In the second condition (denaturing condition) the pellet resulting from cell lysis was resuspended in PBS buffer with 6M urea, shaken for 2 hours and centrifuged. The supernatant was stored at −20° C. Subsequently, the fraction solubilized with urea is subjected to chromatography on metal affinity columns (IMAC). Unlike hybrid conditions, in denaturing conditions, the protein is eluted with PBS containing 6M urea. with a gradient of pH 7.8; pH 6.0; pH 5.3 and pH 4. The protein is purified at pH 4.0. It was then stored at 4°C until use.

#### SDS-PAGE and Western Blot Assays

The purified protein N was brought to 95° C for 10 min and diluted in 4 times Laemmli sample buffer (4% SDS, 20% glycerol, 10% 2-mercaptoethanol, 0.004% bromophenol blue and 0.125 M Tris HCl, pH 6.8). After electrophoresis, the gels were stained for one hour with Coomassie Brilliant Blue (Coomassie Brilliant Blue R-250 0.1%, acetic acid 10%, methanol 50%) and subsequently discolored with a solution of cold methanol. 50%. In Wester-blot assays, the protein was transferred to Immobilon-P membranes (Milipore, USA) in transfer buffer (Tris 25 mM, glycine 192 mM and methanol 20%) at 2 mA / cm2. for 30 min, using a semi-dry transfer kit (Bio-Rad, USA). The membrane was blocked with skim milk 5%, for 1 h at 4° C, washed three times with PBS-Tween 20 0.05%. In some tests, the membrane was cut into strips to evaluate individual sera. It was incubated with 3 sera positive people tested with the IgG / IgM rapid kit for COVID-19 (Abbott), the dilution of the sera was 1:50. Three other membranes were incubated with 3 healthy sera using the same dilution, in blocking buffer at room temperature (25°C) for 30 min. The membrane was washed with PBS-Tween 20 0.05% and incubated with peroxidase-conjugated anti-human IgG antibody (Sigma, USA) at a dilution of 1: 1000, in blocking buffer for 1 h at 37°C, and finally wash with PBS-Tween 20 0.05%. The protein was detected with a revealing solution of diaminobenzidine-H_2_O_2_. The positive and negative sera were provided by a health center, with prior informed consent to the patients.

## Results and Discussion

### Synthetic construction of nucleocapsid N

The nucleocapsid sequence of SARS-CoV-2 was obtained from accession number MN908947.3, which corresponds to the complete genome of the new coronavirus 2 of the isolated severe acute respiratory syndrome coronavirus 2 Wuhan-Hu-1. From the genome, the coding sequence of the nucleocapsid (region 28274-29533), was selected. The sequence is 1289 bp which code for a theoretical 45.6 kDa protein. In order to express the gene in *Escherichia coli*, the codon bias was changed (Figure 1) and it was obtained by chemical synthesis, and it was cloned in the expression plasmid pET-20b. This vector has an export sequence to the periplasm (pelb) that allows the protein to be purified by osmotic shock and a tag of six histidines at the C-terminal end that allows it to be purified by metal exchange chromatography (IMAC). When comparing the synthetic sequence with the wild one, 67.54% changes in the triplets are observed.

**Figure 1.**
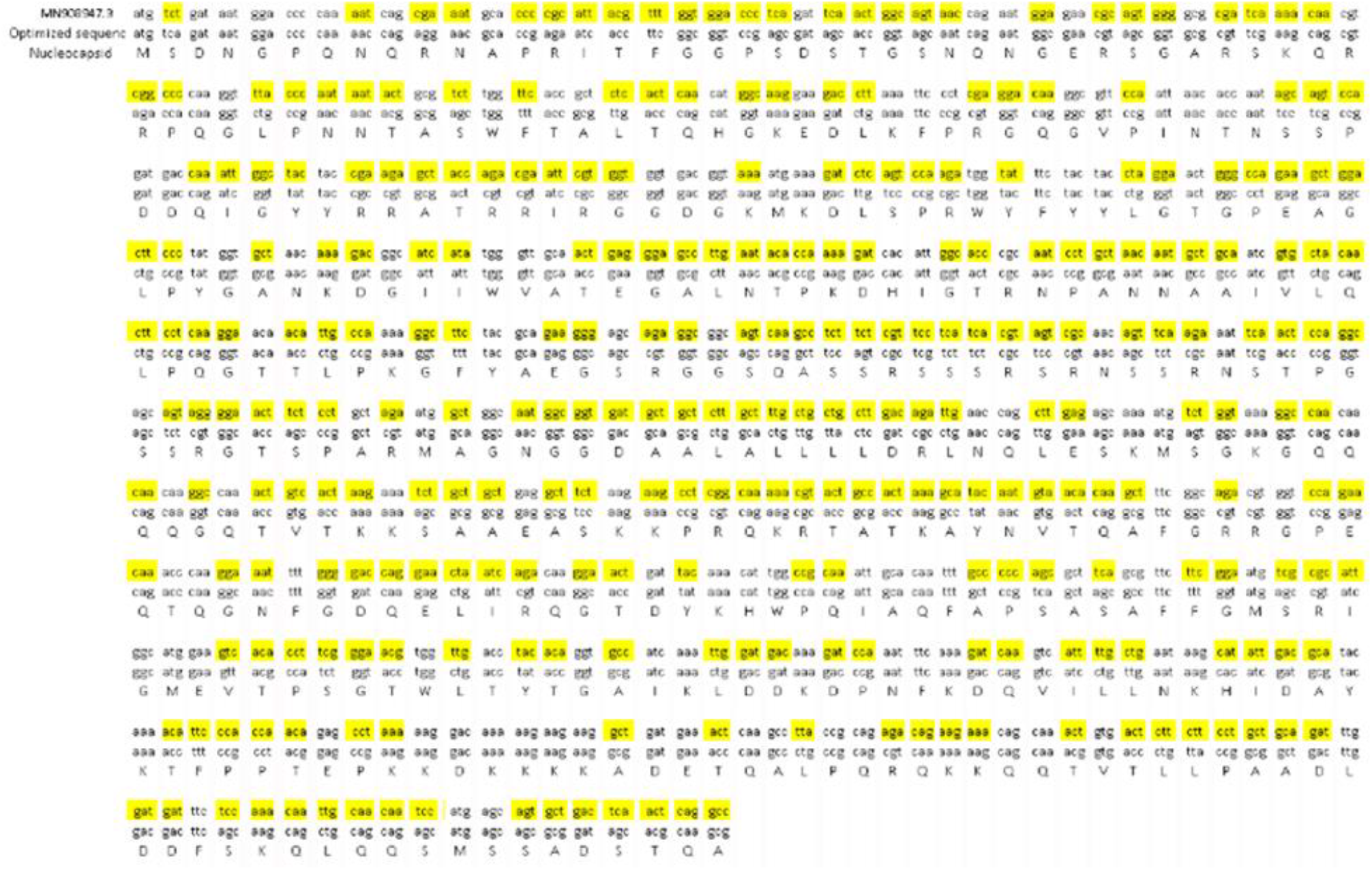
Codon bias of the SARS-CoV-2 nucleocapsid sequence, MN908947.3: 28274-29533. The triplets that were substituted to obtain an optimized sequence to be expressed in Escherichia coli are indicated in yellow. Below the sequences of each codon its corresponding amino acid is indicated.

The plasmid is used as a template to amplify the insert using the forward and reverse oligos of T7, using the following parameters: heating at 95° C for 5 min; 32 cycles of denaturation at 95°C for 1 min, hybridization at 45°C for 3 min and extension at 72°C for 1 min, and a final elongation at 72°C for 5 min, a band of 1610 bp was obtained, (Figure 2), which corresponds to the insert plus part of the vector.

**Figure 2.**
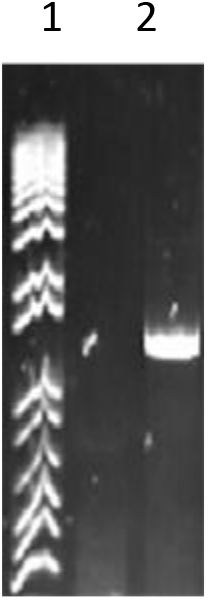
Product of the amplification of the N insert from vector pET20b. Line 1: Molecular weight marker.

### Expression of protein N in *E. coli* Bl21 (DE3)

The *E. coli* strain Bl21 (DE3) was used for recombinant N expression tests. A band with a molecular weight of 50 kDa was obtained, evidenced in 15% SDS-PAGE gels. Protein N was located in the insoluble fraction, forming inclusion bodies, so the osmotic shock strategy was not suitable. The intense band of molecular weight of 15 kDA corresponds to the lysozyme used in the disruption of bacteria (Figure 3).

**Figure 3.**
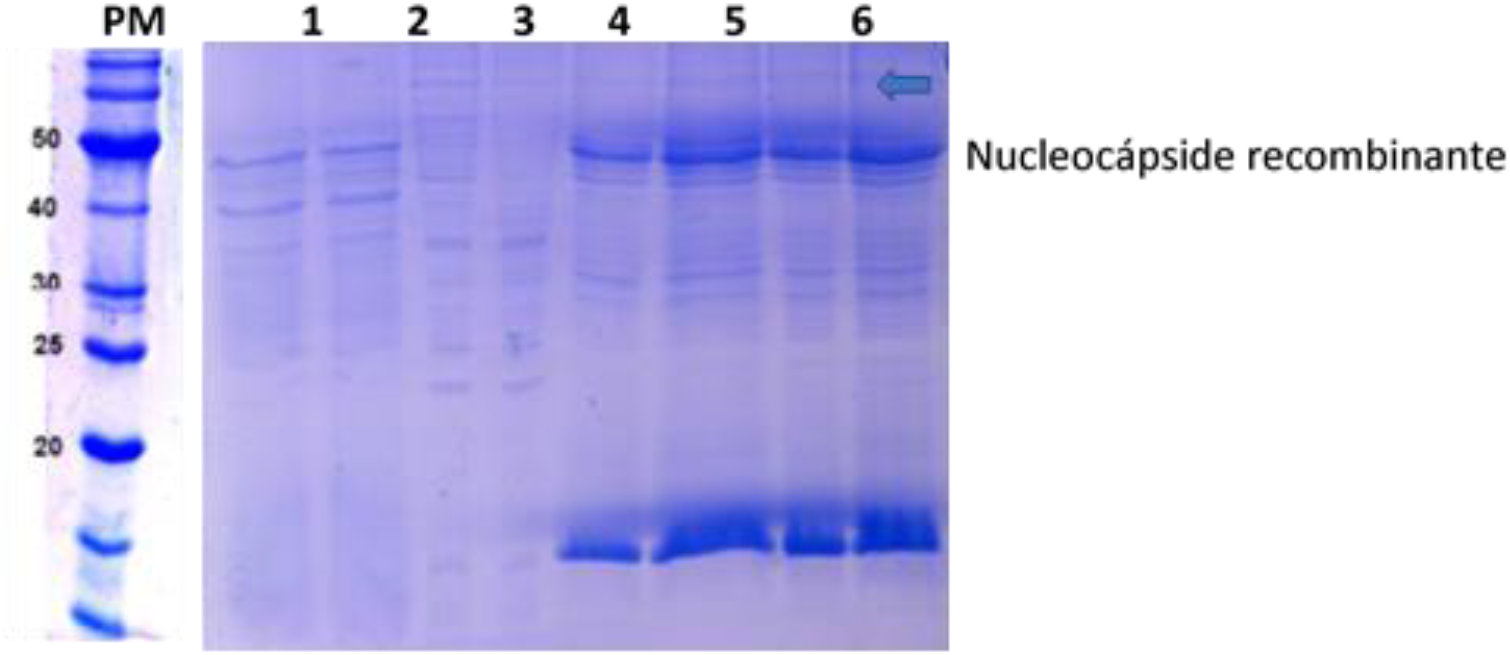
Polyacrylamide gels with S.D.S at 15% of the fractions obtained by inducing the recombinant nucleocapsid with 1 mM IPTG at 37 oC for 16 h. PM: Molecular weight marker. Channel 1-2: Cell extract; Channel 3-4: Soluble fraction. Channels 5-8: Insoluble fraction. 50 ng of protein was used per lane. The apparent molecular weight of recombinant N is 50 kDA.

### Purification and Western-blot of the recombinant nucleocapsid (protein N)

Since it was not possible to use native conditions for purification, due to the nucleocapsid forming inclusion bodies, purification methodologies were carried out under conditions; hybrid and denaturing. The hybrid condition uses guanidine to solubilize the inclusion bodies, after centrifugation, the supernatant is loaded onto the metal affinity column (IMAC), washed with a guanidine buffer to eliminate unfixed proteins and then washed with PBS buffer without guanidine. The protein is eluted with 250 mM imidazole (Figure 4A). loss of protein by degradation is observed and, in addition, when performing the Wester-blot an antigenic protein of *E.coli,* of approximately 60 kDa, is evidenced, which in case of using the ELISA technique would cause false positives (Figure 4C).

**Figure 4.**
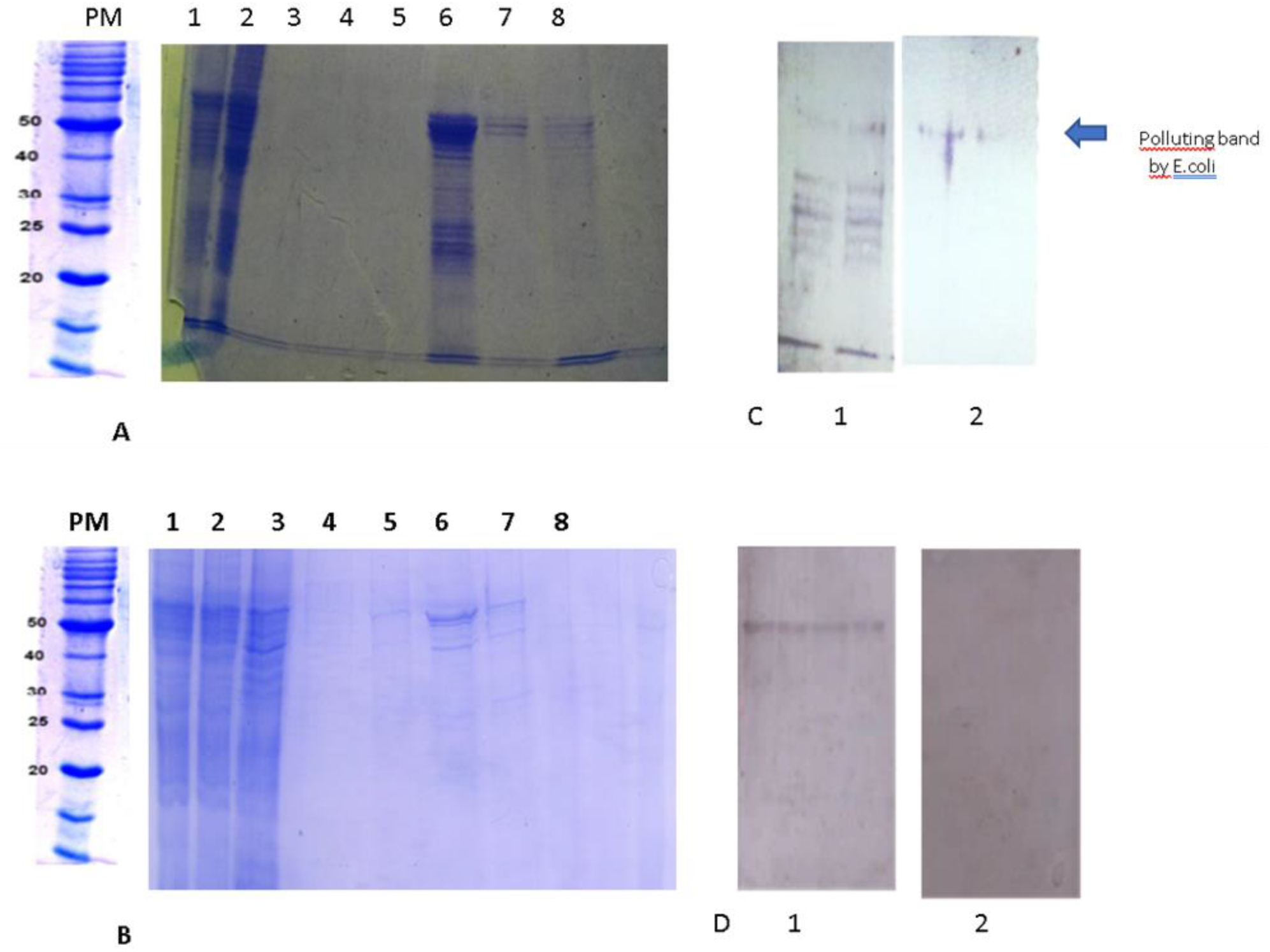
Analysis by SDS-PAGE 12% of the fractions obtained by purifying the recombinant N in: (A). Hybrid condition 1: lane 1: Induced extract. Lane 2: Insoluble fraction. Lane 3: eluted. Lane 4: fraction eluted with imidazole 50 mM, Lane 5: fraction eluted with imidazole 100 mM. Lane 6 fraction eluted with imidazole 250 mM. Lane 7: fraction eluted with imidazole 500 mM. Lane 8: fraction eluted with imidazole 1000 mM. (B). Denaturing condition: Lanes 1 and 2: induced fraction, Lane 3: washing. Lane 4-6: Elusion with, pH 7.0; 6.0 and 5.3. Lane 7: Elusion at pH 4.0, Lane8: elusion at pH 2.0. (C). western blot of positive and negative sera for COVID 19 using the fraction eluted with imidazole 250 mM under hybrid conditions, the samples were placed in duplicate; (D) Western blot of positive and negative sera for COVID 19 using the fraction eluted at pH 5.3 under denaturing conditions, the samples were placed in duplicate. The apparent molecular weight of recombinant N is 50 kDA. PM: Molecular weight marker.

The methodology under denaturing conditions, includes urea in all solutions used to elute the nucleocapsid attached to the metal affinity column (IMAC), the elusion is achieved with a stepped pH gradient. At pH 4.0, the nucleocapsid is eluted with a high degree of homogeneity (Figure 4B).

It is important to note that the fraction obtained under hybrid conditions, exhibited little homogeneity and was unstable, degrading even in the presence of protease inhibitors at −20°C, and it also included *E. coli* contaminants observed in western blots (Figure 4C). When examining the profile of the western blot with the isolated fraction under hybrid conditions, the autoproteolytic degradation recently reported is evidenced (22), due to the bands that appear in the reactions with positive serum, which exhibit different molecular weights and are not achieved. in reaction with COVID-19 negative serum (Figure 4C).

However, unlike purification under hybrid conditions, the denaturing condition yielded a fraction with a high degree of homogeneity (fig 4B), and also with very good stability. Stable solutions of the nucleoprotein are still preserved, even after two months of purification. The nucleoprotein profile is unaltered and reproducible, the reaction exhibited in western blot is highly specific as shown in Figure 4D.

When comparing the previous methodologies; The advantage resulting from the use of the denaturing condition is evident. It is then possible to use this methodology routinely to obtain recombinant nucleoprotein antigen, conveniently useful in diagnostic assays, given the complete absence of reaction with COVID-19 negative sera and its high stability, making it possible to conceive the development of a diagnostic assay. on a large scale such as ELISA kits or rapid tests. The strategy described under denaturing conditions is a simplified purification alternative to that described by De Marco Verissimo C (14) and Djukic et al (19) who report its expression in soluble form and Li (17) where it is obtained by forming bodies of inclusion.

### Wester-blot with positive and negative covid-19 sera

The fraction eluted under denaturing conditions and at pH 5.3 was evaluated with 3 positive sera and 3 negative sera in western blot (5 ug / line), observing reaction bands in the 3 positive sera for COVID-19 (Figure 5A). There was no reaction with the COVID-19 negative sera (Figure 5B). The difference in intensity of the bands may be due to the antibody titers, since these depend on the immunological window period the patient is in.

**Figure 5.**
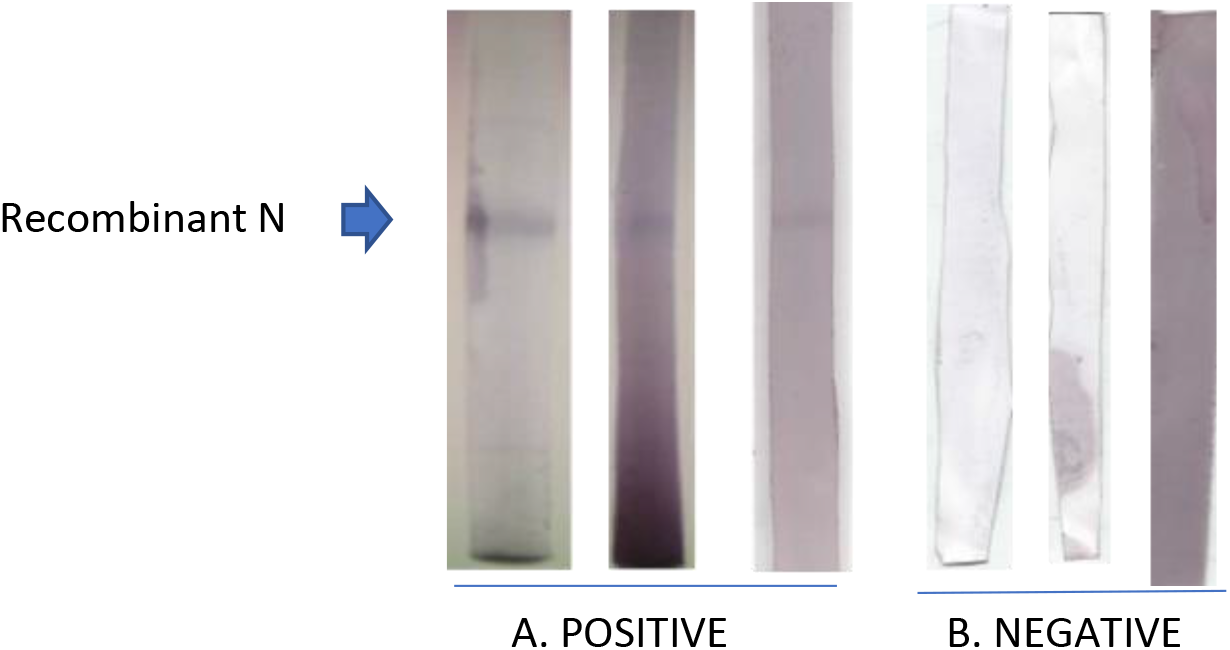
Western-blot showing the presence and absence of antibodies against recombinant N in positive sera A, and negative sera B.

## Conclusion

An alternative method of purification of the recombinant nucleocapsid of SARS-CoV-2, expressed in *E.coli* and using only IMAC and denaturing conditions, is proposed. Unlike most established purification protocols, it does not use strong solubilizing or sonication agents. The procedure is relatively simple and inexpensive and could be carried out on a large scale to obtain an antigen that could be used in the development of serological diagnostic methods for COVID-19.

## Acknowledgment

Mauro Herrera President of CIDCITEI INTERNACIONAL, for the donation of the plasmid pET20b-N.

## Financing

Project 2007001425 FONACIT “Development and / or adaptation of supplies to be applied to kits for serological diagnosis for infectious diseases”

